# Different encoding of reward location in dorsal and ventral hippocampus

**DOI:** 10.1101/2021.09.07.459245

**Authors:** Przemyslaw Jarzebowski, Y. Audrey Hay, Benjamin F. Grewe, Ole Paulsen

## Abstract

Hippocampal neurons encode a cognitive map for spatial navigation^1^. When they fire at specific locations in the environment, they are known as place cells^2^. In the dorsal hippocampus place cells accumulate at current navigational goals, such as learned reward locations^3–6^. In the intermediate-to-ventral hippocampus (here collectively referred to as ventral hippocampus), neurons fire across larger place fields^7–10^ and regulate reward-seeking behavior^11–16^, but little is known about their involvement in reward-directed navigation. Here, we compared the encoding of learned reward locations in the dorsal and ventral hippocampus during spatial navigation. We used calcium imaging with a head-mounted microscope to track the activity of CA1 cells over multiple days during which mice learned different reward locations. In dorsal CA1 (dCA1), the overall number of active place cells increased in anticipation of reward but the recruited cells changed with the reward location. In ventral CA1 (vCA1), the activity of the same cells anticipated the reward locations. Our results support a model in which the dCA1 cognitive map incorporates a changing population of cells to encode reward proximity through increased population activity, while the vCA1 provides a reward-predictive code in the activity of a specific subpopulation of cells. Both of these location-invariant codes persisted over time, and together they provide a dual hippocampal reward-location code, assisting goal- directed navigation^17, 18^.

## Results

To track the activity of the same dCA1 and vCA1 cells when mice learned different reward locations, we imaged over multiple days calcium fluorescence of excitatory cells in Thy1- GCaMP6f mice^19^ with a head-mounted, miniature fluorescent microscope^20, 21^ (Figure 1A–C, S1A). We imaged daily 170 ± 19 dCA1 cells from seven animals and 70 ± 11 vCA1 cells from six animals and matched the identity of active cells between days (Figure S1B). The matched cells had low displacement (mean distance 1.9 µm, IQR: 1.1–3.1 µm) and highly correlated regions of interest (median 0.72, IQR: 0.66–0.77, Figure S4C–D).

**Figure 1.**
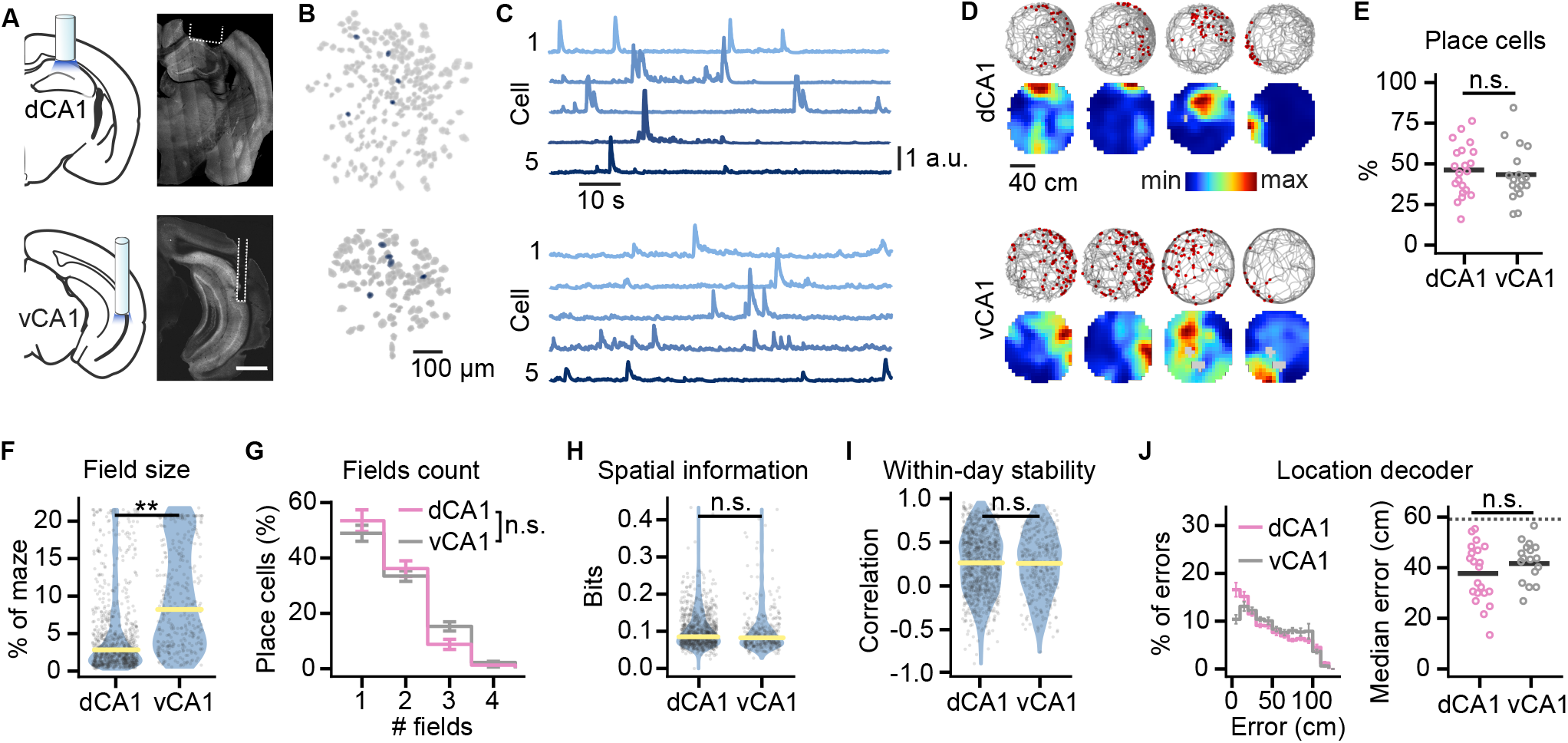
Similar spatial information encoded in dCA1 and vCA1 during foraging. (A) Location of the GRIN lens implanted above dCA1 (top) and vCA1 (bottom) pyramidal cells expressing GCaMP6f. Scale bar 1 mm. (B) Spatial footprints of cells detected in a single day from a dCA1- (top) and vCA1- implanted (bottom) mouse. (C) Example background-subtracted fluorescence traces from the five cells shown in blue in (B). (D) Examples of dCA1 (top) and vCA1 place cells (bottom) recorded during single-day foraging sessions. Locations of calcium events marked with a red dot are overlaid over mouse movement paths (top); place maps are shown below. Gray pixels represent unsampled locations. (E) Percentage of the dCA1 and vCA1 cells identified as place cells during foraging. (F) Field sizes of the dCA1 and vCA1 place cells. (G) Field counts per place cell. (H) Spatial information of place cells normalized by the cell’s mean deconvolved activity. (I) Within-day stability of place fields measured as a correlation between place maps from the same-day early and late trials. (J) Accuracy of decoding location from neural activity in the dCA1 and vCA1. Shows distribution of decoding errors (left) and the median error (right) calculated using cross- validation on single-day activity. Decoders were trained and evaluated on 30 sampled cells to match the cell counts in dCA1 and vCA1; the sampling was repeated 50 times. Distribution of the values shown on violin plots of the width proportional to density; horizontal bars mark the means; individual data points overlayed on top of the violin plots. Error bars mark ± SEM. To avoid double counting of the cells sampled on different days, data in (F) and (H–I) is for place cells recorded in the last-day foraging sessions. The effect of the recording location was tested with linear mixed-effects models in (E), (G) and (J) and with log-linear mixed- effects models in (F) and (H). **p < 0.01.

### dCA1 and vCA1 encoded comparable spatial information but the vCA1 fields were larger

First, to confirm that calcium signals yield similar place cell properties to those previously reported using tetrode recordings, we characterized dCA1 and vCA1 place-specific neuronal responses during foraging on the same maze as that used later during learning of reward locations (Figure 1D, Video S1–2). Because ventral hippocampal lesions increase the mobility of mice^22^, we confirmed that the surgical procedures did not lead to differences in mobility between the dCA1 and vCA1 implanted mice. The two groups were running in a similar fraction of the trials (linear mixed-effects model: F_(1, 11)_ = 10^-4^, p = 0.99; BF_10_ = 0.22; Figure S1E) and with similar speed (log-linear mixed-effects model: F_(1, 11)_ = 0.13, p = 0.73; BF_10_ = 0.28; Figure S1F).

We identified 46 ± 3% of the dCA1 and 43 ± 4% of the vCA1 cells as place cells (see Methods; inconclusive evidence for difference: BF_10_ = 0.45, CI = [-15%, 11%]; Figure 1E; Table S1). In agreement with reports using tetrode recordings in the CA1^7–10^, the vCA1 place fields were larger than dCA1 place fields (strong evidence: BF_10_ = 12, CI = [24%, 204%]; Figure 1F; Table S1). Nearly half of the place cells were active in more than a single location, resulting in multiple place fields. The mean count of place fields per place cell was similar in dCA1 and vCA1 (inconclusive evidence for difference: BF_10_ = 0.80, CI = [-16%, 4%]; Figure 1G; Table S1). Studies using tetrode recordings reported that vCA1 place cells contained less spatial information per spike^7, 8, 10^. Here, their spatial information normalized by the cell‘s mean activity did not differ (moderate evidence for the lack of difference: BF_10_ = 0.25, CI = [-31%, 65%]; Figure 1H; Table S1). However, one-photon calcium imaging could be less sensitive to individual spikes^23^, thus failing to capture the full spatial information. Stability did not differ between dCA1 and vCA1 place fields (moderate evidence for the lack of difference in correlation between same-day early and late trials: BF_10_ = 0.22, CI = [-0.12, 0.19]; Figure 1I; Table S1), and activity of both the dCA1 and vCA1 cell population could be used to decode the animal‘s position with similar accuracy (inconclusive evidence for difference in median decoding error: BF_10_ = 0.8, CI = [-14 cm, 3 cm]; Figure 1J; see Methods).

### dCA1 but not vCA1 place cells accumulated at reward locations

We next compared how the activity of dCA1 and vCA1 place cells changed as a result of learning^3, 4, 6, 10, 24–26^. Mice learned sets of two fixed reward locations in daily sessions. The learning period spanned five days for the first set of locations and two days each for subsequent sets, each with one reward translocated, for a total of three or four sets (Figure 2A). Mice took progressively shorter paths to find the rewards (Figure 2B, Video S3–4). Their memory was tested in unbaited test trials on the day after learning each set (Figure 2C). Mice crossed the reward zones (20-cm-radius disks centered on the learned reward locations) 64 ± 7% more times in the first 120 s of the unbaited test trials compared to the same zones during foraging (linear mixed-effects model, effect of learning: F_(1, 61)_ = 105, p = 10^-14^; BF_10_ = 6 * 10^10^, CI = [45%, 85%]; Figure 2D). Performance of the dCA1 and vCA1 implanted mice did not differ (linear mixed-effects model: F_(1, 16)_ = 0.03, p = 0.87; moderate evidence for the lack of difference: BF_10_ = 0.18, CI = [-13%, 11%], Figure 2D).

**Figure 2.**
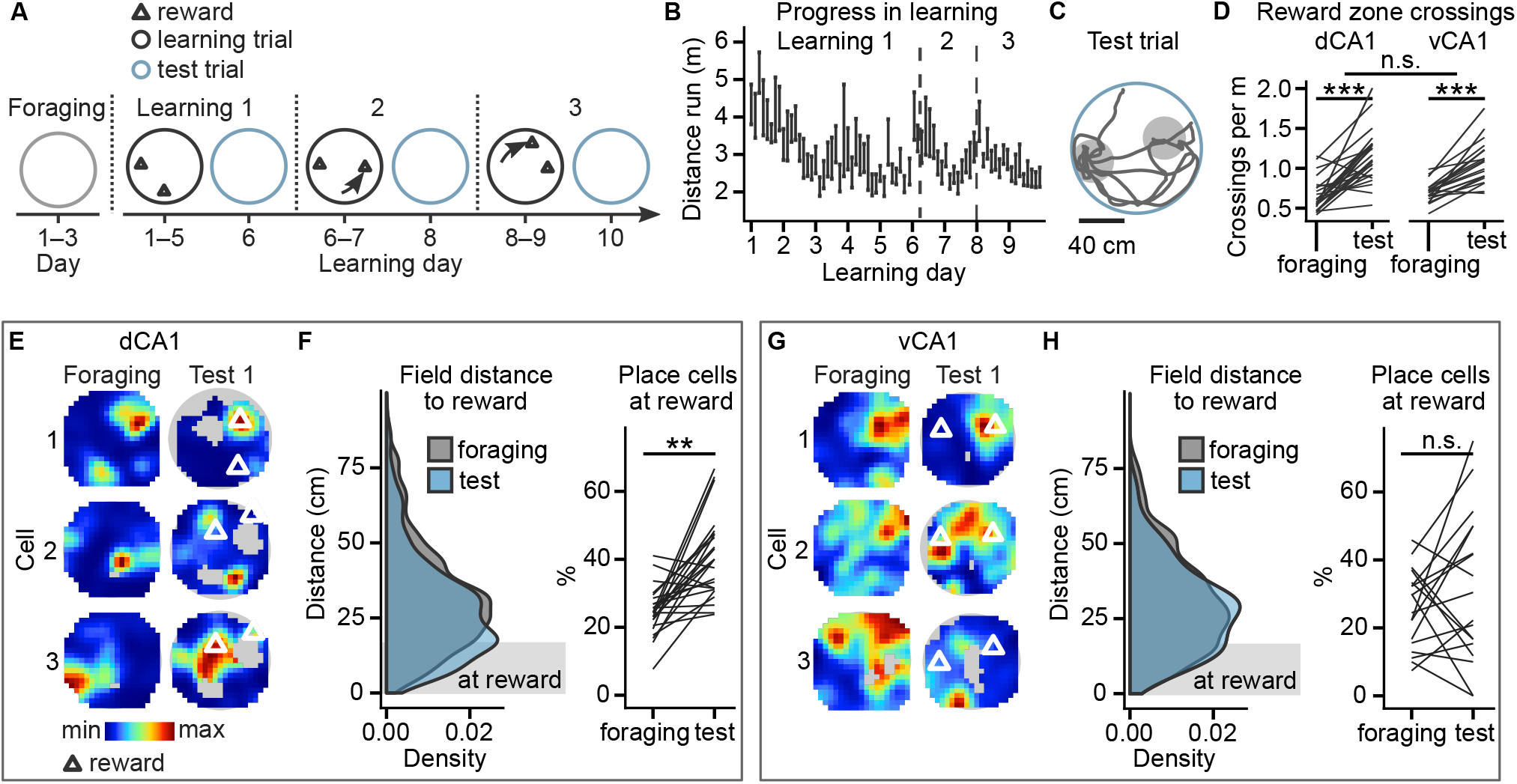
dCA1 but not vCA1 place cells accumulated at learned reward locations. (A) Timeline for the learning and test sessions showing when and how the reward locations (triangles) changed. (B) Progress in learning measured by distance run per trial. Vertical bars mark the mean distance ± SEM, dashed lines mark the time of reward translocations. (C) Example running path of a mouse during unbaited test trial. Gray discs show the extent of the reward zone used for the analysis. (D) Number of reward zone crossings during the first 120 s of the test trials compared to the crossings of the zones centered on the same locations during foraging. Linear mixed- effects model was used to test for the effects of learning, implant location, and their interaction. (E) Examples of dCA1 place fields for the same cell during foraging and the test after Learning 1. Triangles mark reward locations. (F) Learning-induced changes of distance from dCA1 place fields to the closer of two reward locations. Left: distribution of distances shown for place fields from foraging and place fields from unbaited test after learning. Right: the proportion of dCA1 place cells with a place field within 20 cm of the learned reward location (reward field). Data compared with post-hoc test on least-square means of linear mixed-effects model for the effects of learning, implant location, and their interaction. (G) As in (E) but for vCA1 cells. (H) As in (F) but for vCA1 cells. **p < 0.01, ***p < 0.001.

To investigate how the memory of the reward location and the goal-directed behavior affected spatial coding, we compared place fields during unbaited test trials after learning with the foraging trials before learning (Figure 2E,G). The dCA1 but not vCA1 place fields shifted towards the learned reward locations and gained reward fields, defined as fields within 20 cm of one of the reward locations (linear mixed-effects model on the proportion of cells with reward field, trial-type × recording site interaction: F_(1, 13)_ = 16.2, p = 0.001). After learning, the proportion of place cells with a reward field in the dCA1 increased by 65 ± 11% (t_(12.1)_ = 4.8, p = 0.002, BF_10_ = 42850, CI = [34%, 96%]; Figure 2F); whereas it did not change significantly in the vCA1 (inconclusive evidence: t_(13.2)_ = 1.27, p = 0.65, BF_10_ = 0.37, CI = [-18%, 65%]; Figure 2H).

We verified that increased sampling of the reward locations did not account for the increase in dCA1 place cell density. A downsampling procedure that randomly selected an equal count of samples per spatial bin from the foraging and test sessions (see Methods) confirmed the differential effect on place fields in dCA1 and vCA1 (F_(1, 12)_ = 14, p = 0.003; Figure S2). The proportion of place cells with a reward field increased in the dCA1 (t_(1, 11)_ = 4.9, p = 0.002, BF_10_ = 4836, CI = [24%, 71%]), whereas no such change was seen in the vCA1 (moderate evidence for the lack of effect: t_(1, 13)_ = 0.7, p = 1.0, BF_10_ = 0.26, CI = [- 19%, 37%]). It is possible that while the center of mass did not move, the vCA1 place fields could have enlarged towards the rewards. This was not the case, however, as their size decreased from foraging to test trials (log-linear mixed-effects model; F_(1, 973)_ = 35, p = 10^-8^; BF_10_ = 10^6^, CI= [-31%, -17%]).

### dCA1 place cells increased their population activity in anticipation of reward

To gain insight into how memory of reward location affects the population activity, we analyzed dCA1 and vCA1 activity as mice approached the reward. In late learning trials (last day of learning a set of reward locations), the mean dCA1 activity increased by 0.09 ± 0.01 s.d. when mice approached the reward (log-linear mixed-effects model comparing activity at 4–5 s and 0–1 s before the reward, learning stage × reward proximity interaction: F_(2, 1525)_ = 25, p = 10^-11^; late learning increase: t_(1526)_ = 7.8, p = 10^-13^, BF_10_ = 10^11^, CI = [0.08, 0.13] s.d.; Figure 3A,B left, Figure S3A,C). The effect was absent on the first day of learning (early learning, t_(1525)_ = 0.6, p = 1.0, BF_10_ = 0.18, CI = [-0.03, 0.05] s.d.). Also, it was not a direct result of changes in running speed: the effect was absent when the mice stopped at non-rewarded locations (t_(1414)_ = 1.6, p = 1.0, BF_10_ = 0.16, CI = [-0.03, 0.01] s.d.), and it preceded the drop in speed before the reward (Figure S3A). The fraction of active place cells (activity exceeding z-score of 0.5) increased by 33 ± 4%, and of other cells by 15 ± 3% (linear mixed-effects model, cell-type × reward proximity interaction: F_(1, 1085)_ = 10.5, p = 0.001; increase in place cells: t_(1086)_ = 8.0, p = 10^-14^, BF_10_ = 10^10^, CI = [21%, 46%]; increase in other cells: t_(1086)_ = 3.5, p = 0.001, BF_10_ = 168, CI = [6%, 22%]; Figure 3B middle). The increase in the fraction of active place cells was also visible when plotted as a function of distance to the reward (Figure S3E Left), and it correlated with day-mean performance (linear mixed-effects: F_(1, 66)_ = 10, p = 0.002, BF_10_ = 7.2, slope: CI = [0.8, 7.2]; Figure 3C). In contrast, there was no significant correlation of the fraction of active other cells with performance (inconclusive evidence: F_(1, 36)_ = 2.4, p = 0.13; BF_10_ = 0.63, slope CI = [-0.8, 3.1]).

**Figure 3.**
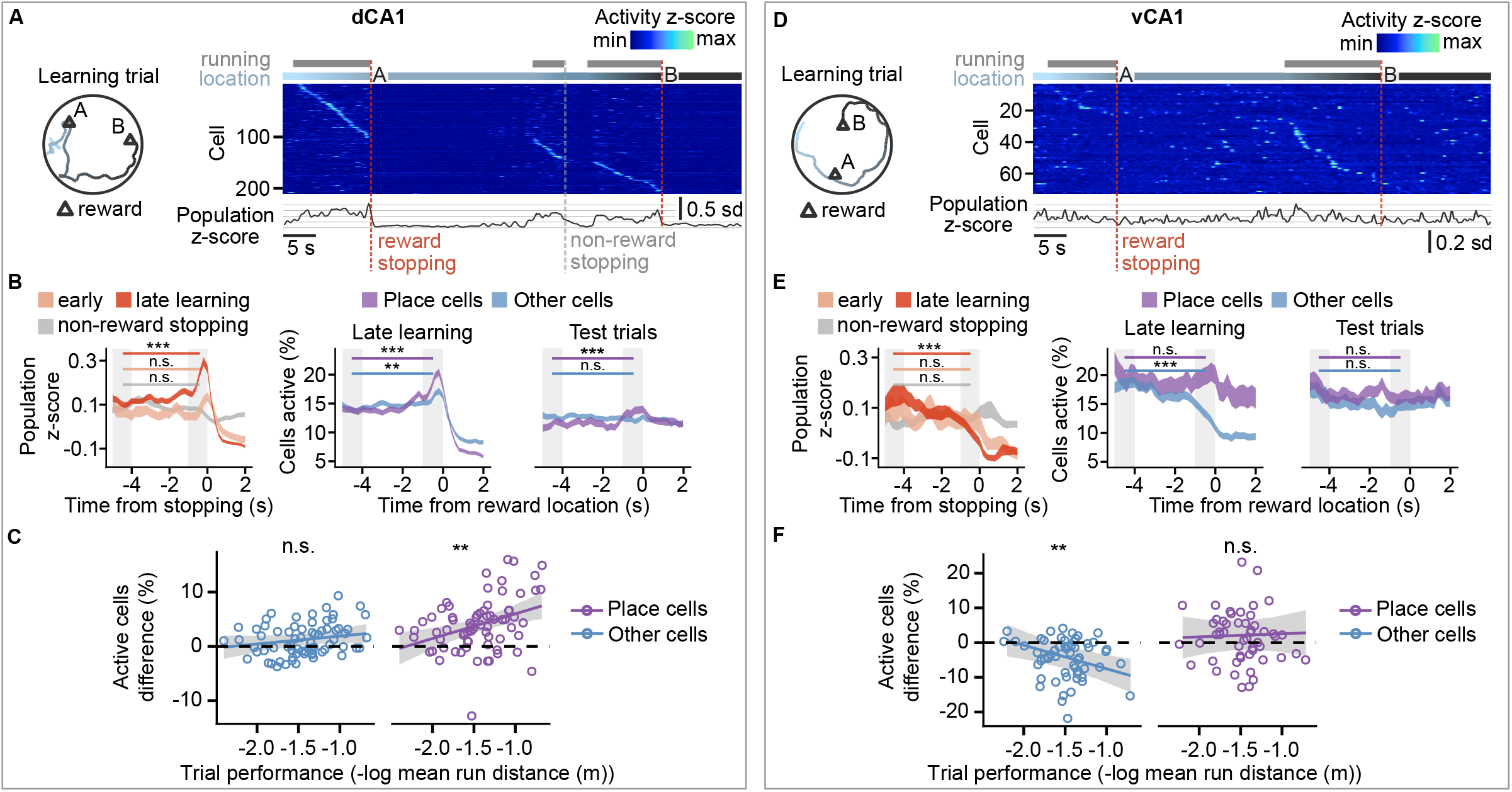
dCA1 population activity ramping up and vCA1 activity ramping down as mice approach the reward. (A) Example single-trial path of a mouse (left) together with dCA1 cell activity (right). Each row of the raster shows the z-scored activity of a single cell. The cells are sorted by the time of their maximum activity. Periods when the mouse ran are marked above the raster with gray. Blue-colored gradient immediately above the raster indicates the color-matched spatial location on the left; the population activity z-score is shown below. Dashed vertical lines show the time when the mouse stopped at rewards or a non-rewarded location. (B) dCA1 population activity as mice approached the reward. After learning, the activity increased before the reward (left), and the percentage of active place cells and non-place cells increased during the approach to reward location (middle). The percentage of active place cells increased during the approach in unbaited test trials (right). The traces have a width of ± SEM; gray rectangles mark 1-s-long periods used for the statistical comparison. Data compared with post-hoc tests on least-square means of linear mixed-effects models for the effects of learning stage, reward proximity, and their interaction. (C) Change in the number of active cells from the 4–5 s before the reward approach to 0–1 s shown as a function of day-mean learning trial performance. The black line shows the slope of modeled regression together with its credibility interval in gray. Linear mixed- effects model used to test for the effects of trial performance. (D) As in (A) but for vCA1. (E) As in (B) but for vCA1. (F) As in (C) but for vCA1. **p < 0.01, ***p < 0.001.

The higher number of active place cells was not caused by reward-associated olfactory cues, as it persisted in the unbaited test trials. When the mice were running the closest to the learned reward location, the fraction of dCA1 active place cells increased by 19 ± 11% while it did not change in other cells (linear mixed-effects model, cell-type × reward location proximity interaction: F_(1, 840)_ = 9, p = 0.003; increase in place cells: t_(840)_ = 3.7, p = 0.001, BF_10_ = 7.4, CI = [6%, 33%]; strong evidence for the lack of change in other cells: t_(85)_ = -0.23, p = 0.82, BF_10_ = 0.09, CI = [-9%, 5%]; Figure 3B right, Figure S3F).

### vCA1 non-place cells decreased their activity in anticipation of reward

Changes in vCA1 population activity contrasted with those in dCA1. In late learning trials, the mean vCA1 activity decreased by 0.09 ± 0.01 s.d. when mice approached the reward (log-linear mixed-effects model, learning stage × reward proximity interaction: F_(2, 1188)_ = 7.3, p = 10^-3^; late learning decrease: t_(1189)_ = -4.2, p = 10^-4^, BF_10_ = 39, CI = [-0.15, -0.4] s.d.; Figure 3D,E left, Figure S3B–C). This effect was absent in early learning trials (t_(1183)_ = 0.82, p = 1.0, BF_10_ = 0.21, CI = [-0.11, 0.5] s.d.) and when mice stopped at non-rewarded locations (t_(1186)_ = -1.0, p = 1.0, BF_10_ = 0.11, CI = [-0.02, 0.05]). The fraction of active place cells did not change but the fraction of active other cells decreased by 32 ± 3% (linear mixed-effects model cell-type × reward proximity interaction: F_(1, 865)_ = 8.8, p = 0.003; no change in place cells: t_(865)_ = 0.3, p = 0.98, BF_10_ = 0.08, CI = [-15%, 23%]; decrease in other cells: t_(865)_ = 4.0, p = 10^-4^, BF_10_ = 10^6^ , CI = [-44%, -21%]; Figure 3E middle, Figure S3D). The decrease was also visible when plotted as a function of distance to the reward (Figure S3E right), and it correlated with day-mean performance (linear mixed-effects: F_(1,55)_ = 8, p = 0.005, BF_10_ = 7.0, slope CI = [-10.5, -1.2]; Figure 3F), whereas there was no correlation between the fraction of active vCA1 place cells and performance (linear mixed-effects: F_(1, 49)_ = 0.9, p = 0.77; BF_10_ = 0.31, slope CI = [-5.4, 6.6]).

In the unbaited test trials, the fraction of active vCA1 place cells and other cells did not change when mice approached learned reward locations (linear mixed-effects model cell-type × reward location proximity interaction: F_(1, 859)_ = 0.7 p = 0.41, reward location effect: F_(1, 859)_ = 1.9, p = 0.16; strong evidence for the absence of change in place cells: BF_10_ = 0.08, CI = [-18%, 12%]; moderate evidence for the absence of change in other cells: BF_10_ = 0.24, CI = [-26%, 3%]; Figure 3E right, Figure S3F). Because the decrease in the fraction of active other cells correlated with the performance only in the baited trials, it is likely that it was related to the reward-associated olfactory stimulus.

### A subpopulation of vCA1 but not dCA1 cells tracked reward location

To investigate whether some cells signaled location-independent anticipation of reward^5, 27^, we compared their activity between test trials performed after the mice learned different reward locations. Of the cells active on the first test trial, 60 ± 6% were active again on the second and 50 ± 5% on the third test trial (Figure S4A). We followed the remapping of place fields between subsequent test trials (Figure 4A). Of the 89 dCA1 place cells with a reward field at the previous reward location, 25% retained their place field, and 31% remapped to either of the current reward locations. However, their place fields were not closer to the current reward locations than those of cells previously without a reward field (log-linear mixed-effects model comparing distances to the closer reward: F_(1, 378)_ = 0.94, p = 0.33, moderate evidence for the lack of difference : BF_10_ = 0.16, CI = [-6 cm, 2 cm]; Figure 4B left).

**Figure 4.**
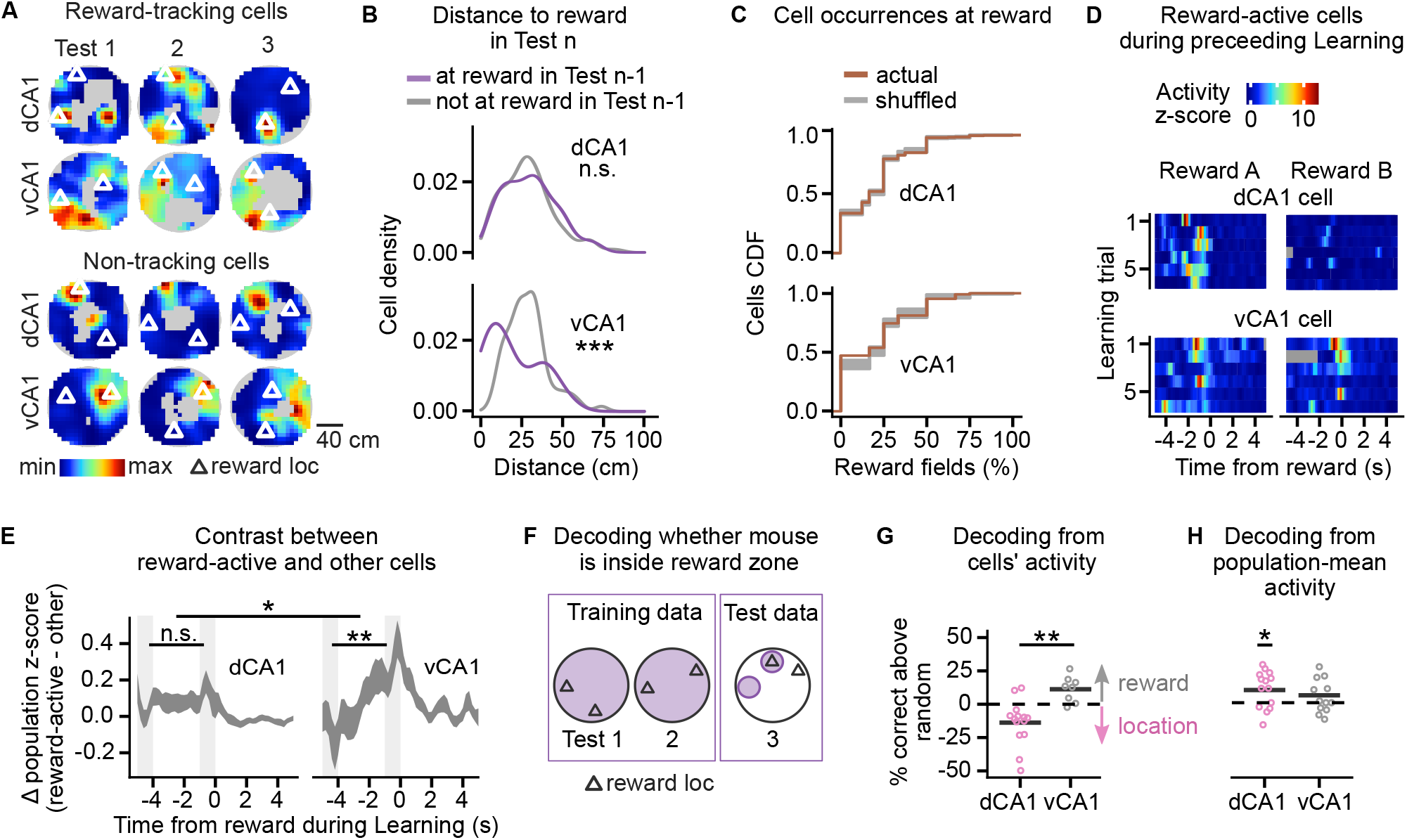
A subpopulation of vCA1 but not dCA1 cells tracked the learned reward location and was active in anticipation of the reward. (A) Example place maps for dCA1 and vCA1 cells with a reward field during Test 1. After the reward location was moved, the reward-tracking cells (top) remapped to one of the current reward locations in Test 2 and 3. The non-tracking cells (bottom) did not remap to one of the current reward locations. (B) Distance from the place field center of mass to the closer of the learned reward locations. Compares the distances between two groups of place cells: (1) cells with a reward field at the previous reward location in the preceeding test trial; (2) cells without a a reward field in the preceeding test trial. Data compared with log-linear mixed-effects model. (C) Cumulative distribution function for the frequency with which cells had a reward field during the test trials. For example, a cell with fields at two reward locations during the three trials had a frequency of 33%. (D) Activity in single-day learning trials of example dCA1 cell and vCA1 cell that had a reward field in the next-day test trial. Each row shows activity in a single trial centered on the time of the mouse arriving at reward. Gray marks periods from before the recording start or after its finish. (E) Population activity difference between reward-active and other cells. Cells were classified as reward-active and other depending on their activity in the next-day test trial. Linear mixed-effects model for the effects of reward proximity, recording site and their interaction; effect of reward proximity in dCA1 and vCA1 tested with least-square means. Gray rectangles mark 1-s-long periods used for the statistical comparison. (F) Training and test data used for binary decoders predicting whether the mouse was running inside reward zone. The decoders were trained on the activity from test trials on two different days. They were tested on activity from another day when the decoder had to flip its prediction for the two tested locations: the previously rewarded location was unrewarded and vice versa. (G) Accuracy of decoding from cells‘ activity shown as the difference from random predictions based on reward zone occupancy probability. Decoders evaluated on data from test trials with different reward locations than in training. For the exact location, the decoder had to give the opposite answer to the training data. Accuracy below the random level means the decoder predicted location rather than predicting reward zone. (H) Shows decoding accuracy from the mean population activity. Accuracy shown as the difference from random prediction. Individual points in (I) and (J) show decoding accuracy per test trial. *p < 0.05, **p < 0.01, ***p < 0.001.

In contrast, the place fields of the vCA1 cells that had a reward field at the previous reward location were subsequently closer to the current reward locations than those of cells previously without a reward field (log-linear mixed-effects model comparing distances from place field to the closer of the two rewards: F_(1, 90)_ = 17, p = 0.0001; strong evidence: BF_10_ = 122, CI = [6 cm, 19 cm]; Figure 4B right). The effect was not due to different place field sizes in the two groups of vCA1 place cells (linear mixed-effects model for mean field size: F_(1, 89)_ = 1.2, p = 0.29; moderate evidence for the lack of difference BF_10_ = 0.32, CI = [- 2.5%, 1.0%]).

The subpopulations of vCA1 place cells with zero or multiple reward fields were larger than expected by chance, suggesting they form two distinct subpopulations – one remapped avoiding and the other remapped tracking reward locations. Of 106 vCA1 place cells, 47.1% had zero reward fields, and 4.7% had reward fields at more than half of reward locations, exceeding the count in 99% and 96% of 1000 cell identity shuffles, respectively (generated by shuffling cell identities assigned to test trial place maps and summing cells‘ reward fields; Figure 4E right). In comparison, of 423 dCA1 place cells, 33.8% had zero reward fields, and 2.1% had reward fields at more than half of reward locations – fractions similar to the respective 34.1% and 2.5% of cells expected by chance (Figure 4E left).

To investigate whether cells with reward fields during test trials were active in anticipation of reward, we analyzed their activity as the mice approached either of the two reward locations during the previous learning day. dCA1 and vCA1 reward-active cells from test trials had higher activity than the other cells as the mice approached reward (log-linear mixed effect models on activity -1–0 s from the reward, dCA1: F_(1, 2134)_ = 20, p = 10^-5^; BF_10_ = 498, CI = [0.1, 0.2] s.d.; vCA1: F_(1, 591)_ = 19, p = 10^-5^; BF_10_ = 224, CI = [0.1, 0.4] s.d.; Figure 4D, S4B–C). The difference between the population activity of the reward-active cells and the other cells increased as mice approached the reward in vCA1 but not in dCA1, suggesting that reward-active cells in vCA1 form a distinct cell population (log-linear mixed-effects model comparing the difference at 4–5 s and 0–1 s before reward, recording site × reward proximity interaction: F_(1, 456)_ = 5.9, p = 0.016; strong evidence for no change in dCA1: t_(455)_ = -0.2, p = 0.90, BF_10_ = 0.10, CI = [-0.1, 0.1] s.d.; strong evidence for change in vCA1: t_(456)_ = -3.3, p = 0.006, BF_10_ = 21.2, CI = [0.1, 0.4] s.d.).

### The vCA1 cell activity and dCA1 population-mean activity predicted reward location with stable and location-independent code

Finally, we wanted to assess to what extent the hippocampus encodes proximity to remembered reward irrespective of its location. We found that hippocampal activity predicted learned reward locations using a code that was shared across different reward locations. To demonstrate that, we first created a binary decoder predicting from dCA1 or vCA1 cell population activity whether the mice were running inside a reward zone during unbaited test trials (Figure 4F). When tested on the same training dataset, decoding from dCA1 and vCA1 had accuracies of respectively 31 ± 2% and 30 ± 3% above that of random predictions based on reward zone occupancy probability (Figure S4D). To show that the activity generalizes to changed reward locations, we evaluated the decoders at one new and one of the previous reward locations from another test trial (Figure 4F).

Decoding from the vCA1 had an accuracy of 10 ± 3% above that of the random predictions. This was significantly higher than decoding from the dCA1 which had an accuracy of 11 ± 4% below that of the random predictions (F_(1, 20)_ = 12, p = 0.002, BF_10_ = 7.1, CI = [3%, 33%]; Figure 4G). The dCA1 decoder gave the same prediction as in the training data even though the reward location was moved, which means it decoded the mouse location rather than reward proximity.

However, because the number of active dCA1 cells ramped up when mice approached the learned reward location (Figure 3A–C), we tested another decoder based on the dCA1 population-mean activity. The decoder used two inputs representing the population activity: the fraction of active place cells and the fraction of active non-place cells. Decoding performed with an accuracy of 10 ± 4% above chance (t_(13)_ = 2.7, p = 0.02, BF_10_ = 3, CI = [1%, 16%]; Figure 4I). Thus, the ramping up of dCA1 activity encoded reward proximity while allowing place cells to encode spatial location.

## Discussion

We found that both dCA1 and vCA1 activity predicts reward location; however, they do so using different codes: (1) The dCA1 anticipated the reward with increased population activity the strength of which correlated with learning performance. The activity engaged changing place cells, allowing independent reward and spatial coding. (2) In vCA1 the same cells were active in anticipation of the reward. No cell was active at all rewards, but their population provided a code for learned reward location that persisted across reward locations and time.

### Comparison of reward location coding in the dCA1 and vCA1

Consistent with previous reports, dCA1 place cells accumulated at the learned reward locations^3, 4, 6, 25, 26^ (Figure 2F) and, as mice approached the reward, the number of active place cells ramped up (Figure 3B). Ramping dCA1 activity was previously reported during reward anticipation in immobile animals^28, 29^. Therefore, it can predict the reward location independently of the spatial representations encoded during movement.

Our findings suggest that the dCA1 place cells that are active at reward locations are part of a flexible place code network rather than a fixed population dedicated to signaling reward locations. We found evidence against the hypothesis that dCA1 cells with place fields close to reward remapped to track the translocated reward better than the other cells (Figure 4D). Thus, rather than a set of neurons specialized for encoding reward locations^5^, a random subset of place cells accumulating at reward locations accounted for the total number of observed place fields at reward per dCA1 place cell (Figure 4E). This evidence suggests that cells are attracted to the reward stochastically, although with probabilities that might differ between them^30–32^.

In contrast, the distribution of vCA1 place fields was unaffected by the memory of reward location. In a study where mice alternated between two marked reward locations, the intermediate CA1 place cells accumulated at the reward locations and were sensitive to the reward value^10^. Possibly, the intermediate CA1 place cells accumulate at reward during a stereotypic running; or in some form of value association. Heterogeneity among vCA1 cells^27, 33^ could be another factor contributing to the difference. In our study, while the non-place cells decreased their activity as the mice approached the learned reward location, a subpopulation of place cells increased their activity and remapped to track the changing reward locations (Figure 3E, 4D–E), similar to the specialized goal-encoding cells that have been suggested to exist in dCA1^5^.

### Function of reward-predictive encoding

Both dCA1 and vCA1 predicted learned reward location using time and location invariant codes (Figure 4G–I). Such population-wide representations might be more reliable than single-function goal cells. Their signal might direct the animal during navigation by increasing their activity in the proximity of a goal^17^, or by signaling reward expectation^18^.

The different encoding of reward-anticipation in dCA1 and vCA1 affects how the signal can be relayed downstream. vCA1 neurons have divergent outputs^33^. The reward-anticipatory subpopulation could target neurons controlling expression of appetitive memory^11, 12, 14, 34–36^, while avoiding those controlling aversion or fear^27, 35, 37, 38^. In dCA1, the ramping-up of population activity in reward-anticipation resembles that seen in the dopaminergic system^39^. Such signal could indiscriminately excite the downstream targets of the dCA1, including nucleus accumbens-projecting neurons that enable conditioned place preference^40^.

Further studies will be required to determine how the reward-anticipatory signals in dCA1 and vCA1 affect activity downstream of the hippocampus. The hippocampal reward- predictive signals could be important for learning and choosing appropriate actions during reward-guided navigation as they are in reinforcement learning models^18, 41^.

## Methods

### Animals

Thirteen adult male Thy1 - GCaMP6f transgenic mice were used in this study^19^ (Jax: 024276). Mice were housed with 2-4 cage-mates in cages with running wheels in a 12:12 h reverse light cycle. All animal experiments were performed under the Animals (Scientific Procedures) Act 1986 Amendment Regulations 2012 following ethical review by the University of Cambridge Animal Welfare and Ethical Review Body (AWERB) under personal and project licenses held by the authors.

### Surgery

Mice underwent two surgeries: the first one to implant a GRIN lens directly above the cells of interest, and the other to fix an aluminum baseplate above the GRIN lens for later attachment of the miniature microscope. The procedures followed the protocol as described in ref ^42^.

Surgeries were carried out following minimal standard for aseptic surgery.

Meloxicam (2 mg.kg^-1^ intraperitoneal) was administered as analgesic 30 min prior to surgery initiation. Mice were anesthetized with isoflurane (5% induction, 1-2% maintenance, Abbott Ltd, Maidenhead, UK) mixed with oxygen as carrier gas (flow rate 1.0-2.0 L.min^-1^) and placed in a stereotaxic frame (David Kopf Instruments, Tujunga, CA, USA). The skull was exposed after skin incision and Bregma and Lambda were aligned horizontally. A craniotomy was drilled above the implantation site. For the dCA1, the craniotomy was 1.5–2 mm in diameter. The cortical tissue and 2 layers of corpus callosum fibers above the hippocampal implantation site were aspirated. Saline was applied throughout the aspiration to prevent desiccation of the tissue. A GRIN lens (1 mm diameter, 4.3 mm length, 0.4 pitch, 0.50 numerical aperture, Grintech) was stereotaxically lowered at coordinates -1.75 AP, 1.75 ML, 1.35–1.40 DV (in mm from Bregma) and fixed to the skull surface with ultraviolet-light curable glue (Loctite 4305) and further fixed with dental adhesive (Metabond, Sun Medical) and dental acrylic cement (Simplex Rapid, Kemdent). A metal head bar was attached to the cranium using dental acrylic cement for head-fixing the animal during the microscope mounting. For the vCA1 implanted mice, a 0.9 mm diameter hole was drilled, and no tissue was aspirated. The GRIN lens (0.6 mm diameter, 4.95 mm length, 1.0 pitch, 0.5 numerical aperture, Grintech) was lowered inside a 21 gauge needle using a custom-made stereotaxic guide that allowed a precise placement of the lens. The lens was placed at coordinates -3.16 AP, 3.6–3.8 ML, 3.40–3.70 DV and the needle guide was retracted allowing for fixation of the lens to the skull surface. After the surgery, the mice were monitored daily for 5 days and given oral Meloxicam as analgesic.

If the GCaMP6f expression was visible in the implanted mice, 4 weeks later the animals were anesthetized for the purpose of attaching a baseplate for the microscope above the top of the GRIN lens. The baseplate was cemented into place and the miniscope was unlocked and detached from the baseplate.

### Histological processing

Following the behavioral experiments, animals were terminally anesthetized by intra- peritoneal injection of pentobarbital (533 mg.kg^-1^) and then transcardially perfused with phosphate-buffered saline (PBS) followed by 4% paraformaldehyde (PFA). Brains were removed and post-fixed for 24–48 hours, then rinsed and subsequently cryoprotected overnight in 30% (w/v) sucrose dissolved in phosphate-buffered saline (PBS). Coronal sections of the hippocampus were cut using a microtome (Leica) with 80–100 μm m thickness.

After rinsing in PBS, the sections were mounted in Fluoroshield with DAPI (Sigma). Sections were examined with a Leica Microsystems SP8 confocal microscope using the 10× and 20× magnification objectives.

### Cheeseboard maze task

The mice performed a rewarded spatial navigation task on a 120 cm diameter cheeseboard maze^3^ with 177 evenly spaced wells. The rewarded wells were baited with ∼100 μm L of condensed milk mixed 1:1 with water.

For the first three days, the mice foraged for rewards baited in randomly selected wells. The mice explored the cheeseboard in three or four trials for a total of 30 minutes per day. A different, random set of wells was baited in each trial.

Next, we performed a spatial learning task. The mice had to learn two locations with baited wells. The baited wells had fixed locations that were at least 40 cm apart chosen pseudorandomly for each mouse. Mice started the trial in one of three locations on the maze: south, east or west. The maze was rotated and wiped with a disinfectant (Dettol) in- between the trials to discourage the use of intra-maze cues. Landmarks of black and white cues were installed on the walls surrounding the maze. The trials were terminated once the mice had consumed both rewards or after 300 s, whichever was sooner. Each learning day consisted of 8 trials with 2–4-minute-long breaks between the trials.

After the first 5-day-long learning period, memory retention was tested on the next day in a 4-to-5-minute-long unbaited trial. The trial was started from a previously unused starting position (north). The performance was measured by the number of reward zone crossings counted when the mouse crossed a circular zone within 20 cm from either of the reward locations. The number of crossings was normalised by the total travelled distance.

Following the learning sessions and memory retention test for the first set of locations, one of the two reward locations was translocated. The new location was pseudorandomly chosen to be at least 40 cm away from the current and previous reward locations. The learning of the new sets of locations was performed over two days and tested in an unbaited trial the following day as described above.

The trials were recorded with an overhead webcam video camera. The video was recorded at 24 Hz frame rate. The mouse body location was tracked with DeepLabCut software^43^, and custom-written software was written to map the mouse coordinates to relative location on the maze. The extracted tracks were smoothed by applying locally weighted scatterplot smoothing (LOWESS) which used moving average of coordinates in 15 video frames. Periods of running were identified when the running speed smoothed with a moving average 0.5 s window exceeded 4 cm/s.

### Calcium imaging

Calcium imaging was acquired using Miniscope – a head-mounted microscope^20^ (v3 and v4 Miniscope). Microscope emitted blue excitation light (∼470 nm spectral peak) 470 nm spectral peak) whose power was adjusted per each animal. Fluorescence emissions were passed through an emission filter (bandpass filter, 525/50 nm) and collected by CMOS imaging sensor. Before the start of the recording, the mouse was head-fixed on a running wheel to attach the microscope and adjust its focal plane so it matched the field of view from the previous recordings. Afterwards, the mouse with the Miniscope attached was placed in a start box for 3–5 minutes before recording sessions started. The calcium imaging was acquired at 20 Hz, and synchronously started with WebCam camera recording.

### Calcium signal processing

CaImAn software was used to motion-correct any movements between the calcium imaging frames, identify the cells and extract their fluorescence signal from the video recordings^44^. The method for cell and signal detection was based on constrained non-negative matrix factorization^45^ (CNMF-E). CaImAn extracted background-subtracted calcium fluorescence values (F) and deconvolved the signal. The deconvolved signal can be interpreted as a scaled probability of a neuron being active. The calcium imaging videos recorded in the same-day trials were motion corrected to a common template frame and were concatenated. Signal extraction and further processing was performed on the resulting long video, allowing the detection of cells and signals present across the trials. To improve the computational performance, the videos were cropped to a rectangle containing the imaged cells and the video width and height was downsampled by a factor of 2.

The putative cells identified were automatically filtered using CaImAn. The results were visually inspected and the filtering parameters adjusted to exclude non-cell like shapes and traces from the filtered components. The criteria used for the filtering included a threshold for signal to noise ratio of the trace, the minimum and maximum size of the component‘s spatial footprint, threshold for consistency of the spatial footprint at different times of the component‘s activation, and a threshold for component‘s resemblence to a neuronal soma as evaluated by a convolutional neural network provided with CaImAn software.

The identity of cells between the recordings on different days was matched using a registration algorithm implemented in CaImAn. The algorithm aligned the image with spatial footprints of cells from all days to the image from the reference day and matched the cells when their centers of mass were closer than 10 µm.

The deconvolved traces were smoothed in time with a Gaussian kernel (σ = 75 ms). The trace was time binned by averaging the values in 200 ms bins.

For the comparison of dCA1 and vCA1 activity, calcium event rates are reported. A calcium event was detected whenever the cell‘s deconvolved signal crossed 20% of its maximum value.

### Place cell detection and analysis

To assess how spatial locations modulated activity of a cell, we considered periods of running as described above and calculated place maps — mean neural activity per spatial bin. The total activity inside 6 x 6 cm bins was summed from the smoothed deconvolved signal. The mean neural activity in the spatial bin was then calculated as a ratio of the total activity to the total occupancy in the bin after both maps were smoothed across the space using a 2D Gaussian kernel with σ = 12 cm. The place map was filtered to include spatial bins with total occupancy that exceeded 1 s (5 time bins, thresholded on unsmoothed total occupancy).

Spatial information of a cell‘s activity was calculated using the place map values.

Spatial information was defined as^46^:

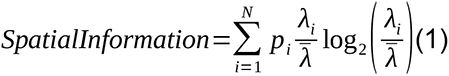

where *λ̄* represents the mean value of the neural signal, *p_i_* represents probability of the occupancy of the i-th bin, and *λ_i_* represents its mean neural activity.

Spatial information was compared to the value expected by chance. The chance level was calculated by circularly shifting the activity with regards to the actual location. For each cell, the activity was circularly shifted within the trial by a time offset chosen randomly (minimum offset 10 s for baited and 20 s for unbaited trials). If the cell‘s spatial information exceeded 95% values calculated on 1000 random shifts of its activity, it was defined as a place cell.

A limited number of neuronal responses sampled per spatial bin can lead to an upward bias in the estimated spatial information^47^. To correct for this bias, we subtracted its estimated value from the estimated spatial information. The bias was estimated as the mean spatial information from the time-shifting procedure used for place cells detection.

This procedure did not require binning the neuronal responses from the calcium imaging as required by analytical estimation^48^, and has been used previously to estimate mutual information bias^49^.

We defined the field size as the fraction of a place map with values exceeding half the maximum value. Fields in the place map were identified by finding local maxima exceeding half the global maximum. The local maxima were restricted to be at least 25 cm apart and have at least one more adjacent spatial bin exceeding half the global maximum. The center of mass for the field was calculated and used to report the field‘s distance from the reward locations. For place cells with multiple place fields, the shortest distance from the reward was used. Fields ≤20 cm from the reward location were referred to as reward fields. To count the reward locations where a place cell had a reward field, only cells that were classified as a place cell in at least half of the test trials were considered.

### Bayesian decoders

Two Bayesian decoders were constructed from the neural activity: the first one decoding spatial location of the running mouse, the second one decoding whether the mouse was running in the proximity of a learned reward location.

The decoders used binarized background-subtracted calcium fluorescence values F. The binarized trace had value 1 (active cell) when the fluorescence exceeded the 90th percentile of the cell‘s values for that day; otherwise, the binarized trace had value 0 (inactive cell).

The Bayesian decoder assumed activity of the cells was independent given the output, and it chose the output to maximize posterior probability given the neural data:

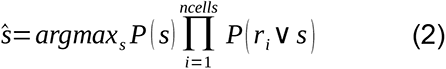

For decoding the mouse location during running, s represents the spatial bin, P(s) represents the prior occupation probability in the spatial bin s, and P(r_i_ | s) represents the probability of the i-th cell being active in the spatial bin s.

For decoding whether the mouse was running in the proximity of learned reward location, s represents if the mouse is within 20 cm from the reward, P(s) represents prior occupation probability, and P(r_i_ | s) represents the probability of the i-th cell being active away or in proximity of the reward.

The decoders were trained and evaluated on two non-overlapping datasets:

(1) The decoder for the spatial location was trained, and evaluated using a cross-validation method as follows: The day‘s session was split into five equal parts. A single part was reserved for evaluation and the others for training the decoder. The decoder was trained and evaluated, and the process was repeated five times, each time with a different part of data reserved for evaluation. The decoder was compared to a baseline random decoder which predicted spatial location based on prior occupancy probabilities. The decoder errors were reported as a distance between the actual and the predicated spatial bin. Because fewer cells were recorded in the vCA1 than in the dCA1, we compared decoders trained on equally sized populations by randomly sampling 30 cells from each recorded session (72% of the vCA1 recordings had more than 30 cells). The spatial decoding from equally sized neuronal populations was repeated 50 times with different cell samples.
(2) The decoder of whether the mouse is at a reward location was trained on data from two unbaited test trials, which were performed on different days and shared a single learned reward location. The training dataset was filtered to times when the mouse was in proximity of the learned reward location (distance ≤15 cm), or the mouse was well-away from the reward location (distance ≥40 cm). The decoder was evaluated on data from another unbaited test trial. In this trial, one of the learned reward locations was different from the ones in the training dataset, and one of the learnt reward locations was missing from the current ones. Only data from the proximity of either of these two locations was used for evaluation (the reward zone vs the previous reward zone). The evaluation was restricted to trials that shared at least 10 cells with the training trials. The decoder assumed equal prior P(s) of the zones. The resulting decoder‘s performance was compared with a baseline random decoder. The decoder errors were reported as the percentage of correct predictions.

### Downsampled data comparison

To verify that differences in maze occupancy between foraging and test trials were not the reason for the observed accumulation of place cells at goal location, we randomly downsampled the data. For each spatial bin in the two sessions, an equally sized subset of timestamps was selected to match the lower of the two occupancies. The selected timestamps were used to construct place maps and to identify place cells. The random downsampling procedure was repeated 100 times, and the statistics about the place field locations and their distance to reward locations aggregated.

### Population activity on reward approach

To analyze the population activity during an approach to reward locations, periods of running that exceeded a minimum duration of 3 s were used. In the baited trials, the running bouts were aligned by the time of the tracked mouse body stopping within 7 cm from the reward. For the bouts stopping at non-rewarded locations, the stops at distance >24 cm from the reward were included. In the unbaited trials, the running bouts were included if they included a location <18 cm from a learned reward location and covered a distance >12 cm. The deconvolved z-scored activity was aligned to the timestamp when the mouse was closest to the learned reward location. The mean population z-scored activity was calculated for 1 s-long bins and the activity at 4–5 s before the bout finish was compared to the activity at 0–1 s.

### Statistical analysis

Results are reported using two statistical methods. First, we estimated p-values using null hypothesis significance testing. The p-values are low for small effects assessed on large sample sizes; they depend on unseen data, and on the plan for how many animals to test experimentally^50^. Therefore, we also report Bayes Factors^51^ — a measure of relative evidence for two competing hypotheses. It is calculated as a ratio of posterior probabilities: the probability of the alternative hypothesis given the observed data over the probability of the null hypothesis given the observed data. We assumed equal prior probability of the alternative and null hypothesis. In addition to providing further statistical support to significant p-values, Bayes Factor analysis gives evidence for the absence of differences where the effects are non-significant^51^. The magnitude of evidence was graded as inconclusive, moderate or strong following Jeffreys‘ thresholds^52^.

Mixed-effects models were used for the statistical analysis to allow for unbalanced sampling and correlated samples. Both apply to this data, for example due to correlations between the samples of cell activity recorded at the same timestamp, or recordings from the same mouse on different trials. The effects were assessed with linear and log-linear mixed-effects models. The fixed effects were the statistically tested effects such as implant location (dCA1 vs vCA1) or cell type (place cell vs non-place cell); the random effects were modeled as mouse-specific and session-specific random variables. The random effects were included in the estimation of the linear regression intercept.

For the frequentist approach, the model coefficients were estimated using the restricted maximum-likelihood method. The residual errors were checked for linear model assumptions: zero mean, no correlation with the predicted values and homoscedasticity. To satisfy these assumptions, some models used a log-linear transformation of the response variable. The significant effects and their interactions were reported and the post-hoc tests were performed on differences in least-square means of the paired groups. The tests used Sattherwaite estimation of degrees of freedom and adjusted p values using Holm-Bonferroni correction.

For the Bayes factor analyses, the mixed-effects models mirrored the frequentist models and had the same fixed and random effects. The priors were specified as cauchy distribution with sqrt(2)/2 scale for fixed effects and 0.5 scale for random effects. These priors follow the expectation that the in-between mice differences are smaller than the effects of interest. Bayes factor for the effect of interest was calculated as probability of the full model over the probability of the model excluding the tested effect.

The effect sizes were reported with 95% credibility intervals (CI; equal-tailed interval). The interval can be interpreted as a range within which the effect falls with 95% probability given the evidence from the observed data. Credibility intervals were estimated from the samples of the model‘s posterior distribution.

Statistical analysis was performed in R version 3.6.3 (ref ^53^). Data are reported as mean ± SEM unless otherwise stated. The linear mixed-effects models were built in R with package ‘lme4’ and p values for the fixed effects were obtained using Sattherwaite estimation of degrees of freedom implemented in the ‘lmerTest’ R package. Least-square means were calculated and tested with ‘lsmeansLT’ function from the same package. Bayesian linear mixed-effects models were created using ‘BayesFactor’ R package and ‘lmBF’ function.

Data are reported as mean ± SEM unless otherwise stated.

### Data and code availability

Code used for the analysis and to generate the figures can be accessed on the authors’ GitHub sites: (1) for processing calcium signal using CaImAn: https://github.com/przemyslawj/caiman_scripts, (2) for package calculation spatial metrics and implementing a Bayesian decoder: https://github.com/przemyslawj/datatrace, (3) for scripts generating the figures in the manuscript: https://github.com/przemyslawj/hpc-reward-coding. Data will be shared on request.

## Supporting information

Video S1. Calcium imaging from dCA1 during foraging.

Video S2. Calcium imaging from vCA1 during foraging.

Video S3. Calcium imaging from dCA1 during learning.

Video S4. Calcium imaging from vCA1 during learning.

## Acknowledgements

We thank Dr Ben Phillips for experimental advice and Dr Julija Krupic and Mr Roman Boehringer for their feedback on the manuscript. We gratefully acknowledge the Cambridge Advanced Imaging Centre for their support and assistance in this work. We also thank Mr Max Schwiening and Dr Christof Schwiening for 3D printing parts of the experimental equipment.

## Author contributions

All authors designed research; P.J. performed research and analyzed data; P.J. wrote the paper with support from B.F.G. and O.P.; all authors approved the final version of the manuscript.

---

**Table S1.** Statistics of dCA1 and vCA1 place cells. Summary of statistics presented in Figure 1.

| Tested variable | mean ± SEM | statistic and p-value | BF<sub>10</sub> | CI | n | Fig |
| --- | --- | --- | --- | --- | --- | --- |
| Place cells fraction | dCA1: 46 ± 3%<br>vCA1: 43 ± 4% | linear mixed-effects<br>$F_{(1,11)} = 0.12$, $p = 0.74$ | 0.45 | [-15%, 11%] | 39 sessions from 13 mice | 1E |
| Place field size (% of maze) | dCA1: 3.4 ± 0.1%<br>vCA1: 8.2 ± 0.4% | log-linear mixed-effects<br>$F_{(1,10)} = 12.5$, $p = 0.006$ | 11.5 | [24%, 204%] | 720 cells from 13 mice | 1F |
| Mean count of fields | dCA1: 1.7 ± 0.04<br>vCA1: 1.6 ± 0.06 | linear mixed-effects<br>$F_{(1,11)} = 1.9$, $p = 0.19$ | 0.8 | [-16%, 4%] | 39 sessions from 13 mice | 1G |
| Normalized spatial information | dCA1: 0.07 ± 2×10<sup>-3</sup><br>vCA1: 0.07 ± 6×10<sup>-3</sup> | log-linear mixed-effects<br>$F_{(1,10)} = 0.15$, $p = 0.71$ | 0.25 | [-31%, 65%] | 720 cells from 13 mice | 1H |
| Place field stability | dCA1: 0.45 ± 0.01<br>vCA1: 0.46 ± 0.02 | linear mixed-effects<br>$F_{(1,10)} = 0.22$, $p = 0.65$ | 0.22 | [-0.12, 0.19] | 720 cells from 13 mice | 1I |

**Video S1. Calcium imaging from dCA1 during foraging.**

Behavior (top left) together with the recorded calcium imaging (top right) and the extracted calcium fluorescence traces for 50 cells (bottom).

**Video S2. Calcium imaging from vCA1 during foraging.**

As in Video S1, but for vCA1.

**Figure S1.**
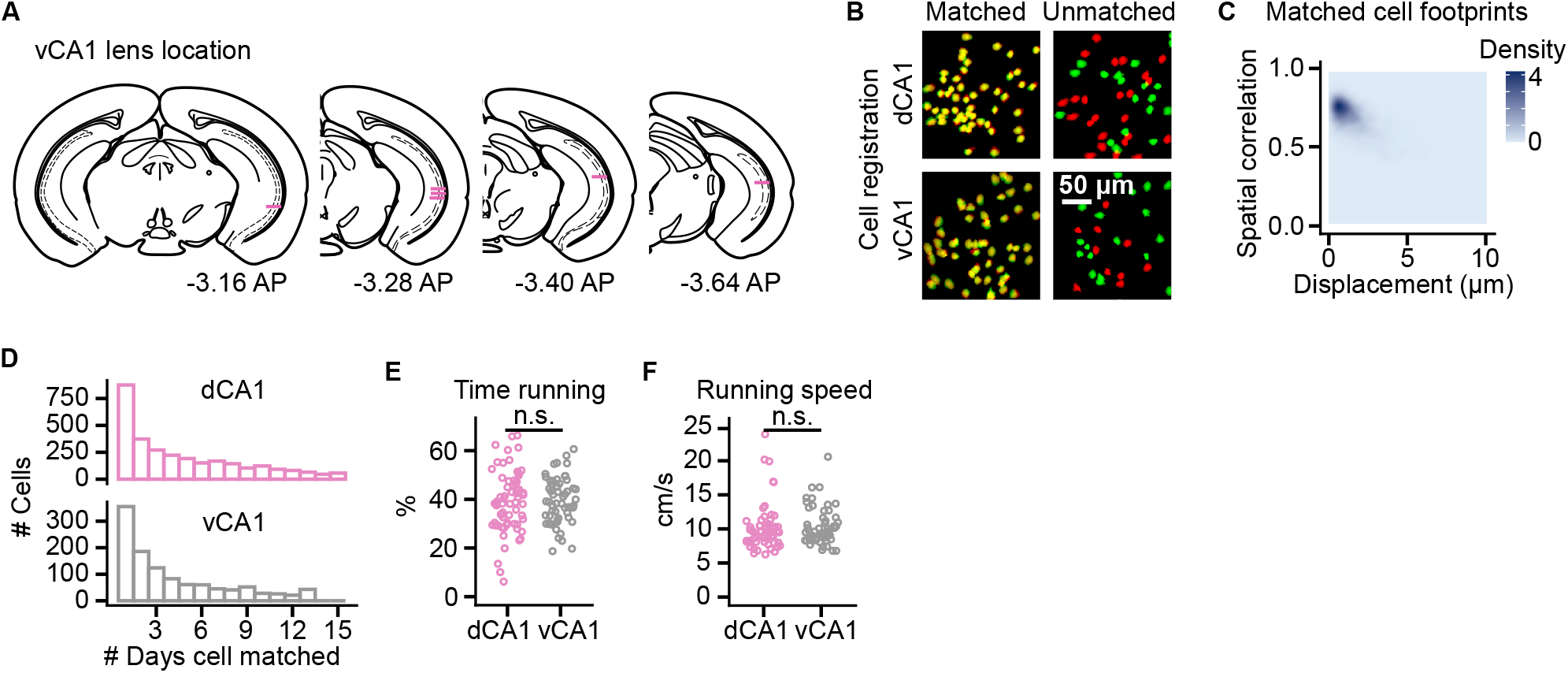
Matching cell identity between days, and modulation of the activity by running, related to Figure 1. (A) Reconstructed location of the recorded cells under the GRIN relay lens implanted in the vCA1 of 6 imaged mice. Horizontal bars mark the bottom of the relay lens. (B) Matching of cell identity (cell registration) based on the cells’ spatial footprints in calcium recordings from two different days. An example with cells found on one day is shown in green, on the other day in red, and their overlap in yellow. (C) Spatial correlation of the matched cells as a function of the distance between their centroids. (D) Histogram for the number of recording days that a cell was active and matched. Over the 14–16 days, there were a total of 2,965 dCA1 and 1,125 vCA1 unique cells. (E) Percentage of cells from the first test trial active again in the later test trials. (F) Percentage of foraging trials that the dCA1 and vCA1 implanted mice spent running. (G) Running speed of the dCA1 and vCA1 implanted mice. The effect of the recording location was tested with linear mixed-effects models in (F) and log-linear mixed-effects models (G).

**Figure S2.**
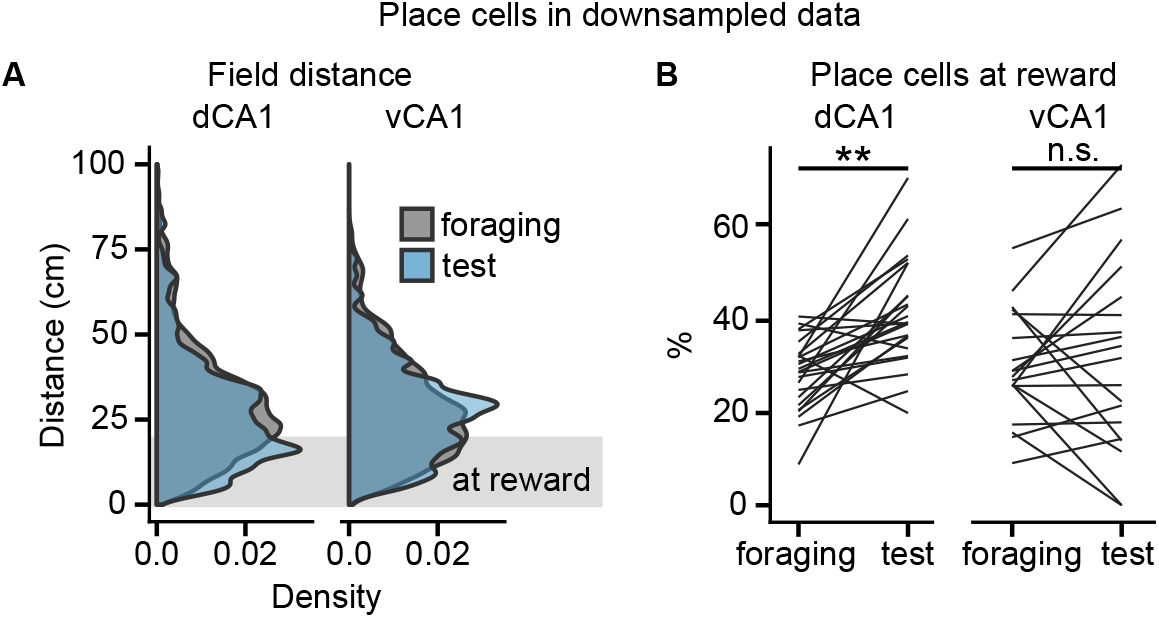
Place cell accumulation at learned reward locations was not caused by increased occupancy at the reward locations. (A) Ss in Figure 2F left and 2H left for place cells calculated on randomly downsampled data to match occupancies between foraging and test trials. (B) Change in the percentage of place cells with a reward field between the foraging and test trials calculated on randomly downsampled data. Data compared with post-hoc test on least-square means of linear mixed-effects model. **p < 0.01.

**Video S3. Calcium imaging from dCA1 during learning.**

Behavior (top left) together with the recorded calcium imaging (top right) and the extracted calcium fluorescence traces for 50 cells (bottom).

**Video S4. Calcium imaging from vCA1 during learning.**

As in Video S3, but for vCA1.

**Figure S3.**
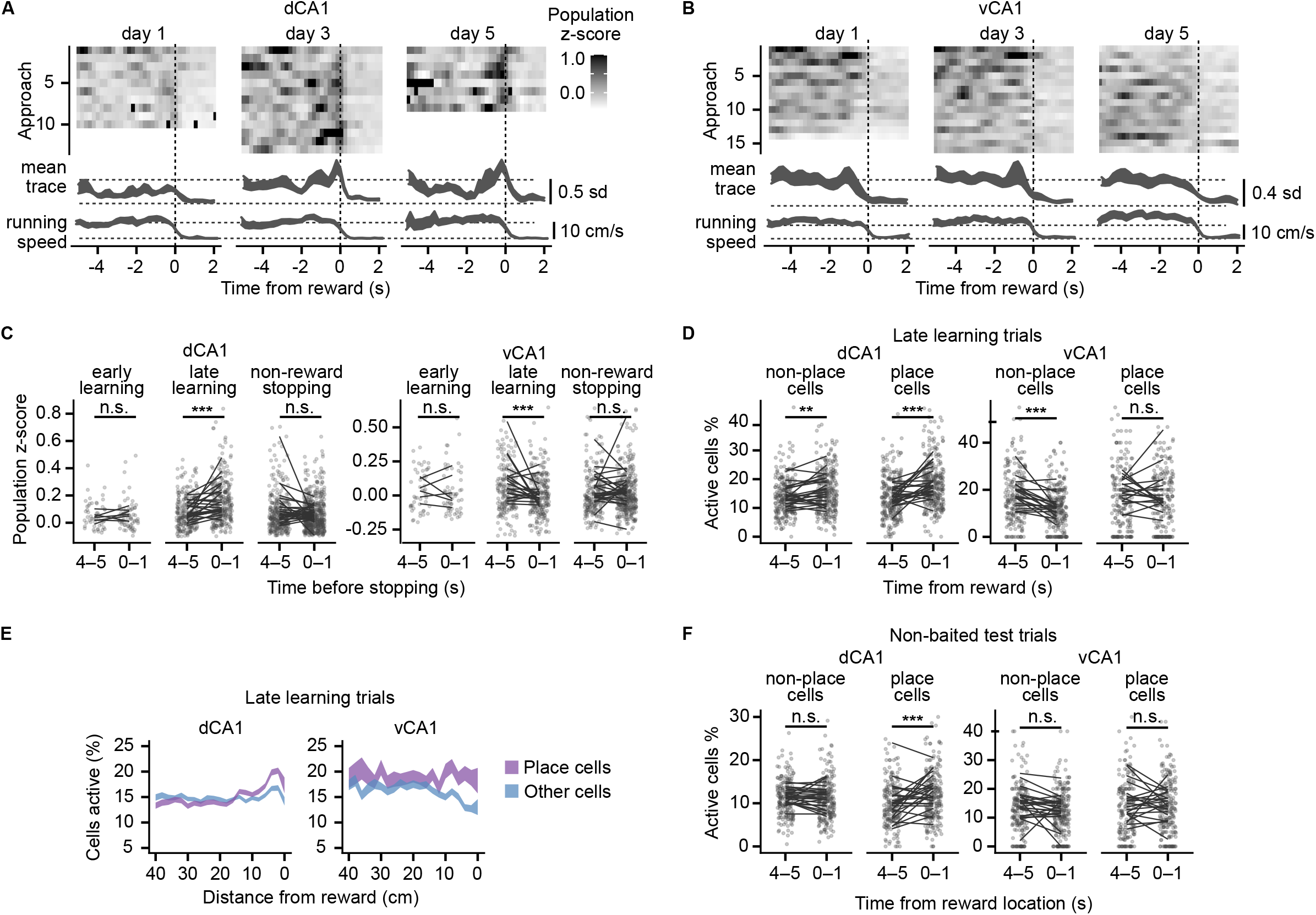
Changes in dCA1 and CA1 population activity, data for individual approaches to the reward in Figure 3. (A) Examples with dCA1 population activity during individual approaches towards the reward locations (middle). Day-mean ± SEM dCA1 activity and running speed shown below the examples. (B) As in (A) but for vCA1. (C) Population activity during 4–5 s and 0–1 s time window before the mouse arrives at the reward. Data points show values for individual running bouts and correspond to the mean ± SEM in 3B and 3E left panels; lines connect day-mean values calculated per animal. (D) As in (C) but for percent of active place cells and the other cells in late learning trials. Data points correspond to the mean ± SEM in 3B and 3E middle panels. (E) Mean population activity during running bouts towards the reward as a function of reward distance. The trace has a width of ± SEM. (F) As in (C) but for averaged percent of active place cells and the other cells in unbaited test trials. Data points correspond to the mean ± SEM in 3B and 3E right panels. Data compared with post-hoc tests on least-square means of linear mixed-effects models for the effects of learning stage, reward proximity and their interaction. **p < 0.01, ***p < 0.001.

**Figure S4.**
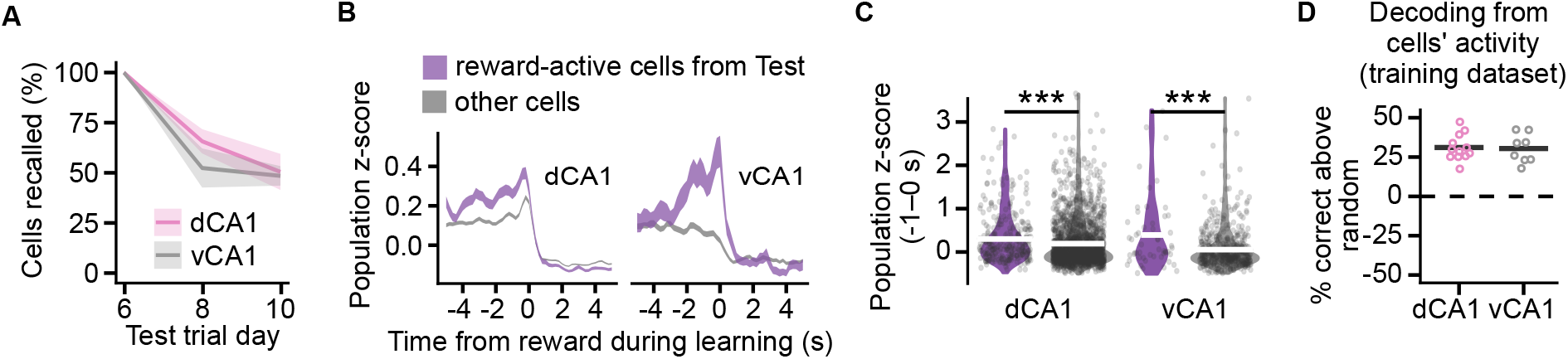
Anticipatory activity of reward-active cells, and decoding reward location, related to Figure 4. (A) Percentage of cells from the first test trial active again in the later test trials. The ribbon has a width of ± SEM. (B) Averaged activity of two cell groups: cells with reward fields in a next-day test trial and the cells without a reward field shown around the time of mice arriving at the reward. The trace has a width of ± SEM. (C) Day-mean activity of cells 0–1 s before the mice approached the rewards during late learning trials. Compares the activity of the cells with and without reward fields during next-day test trials. Distribution of the values is shown on violin plots of the width proportional to density; horizontal bars mark the means. Data compared with post-hoc tests on least-square means of linear mixed-effects models for the effects of cell group (reward active or not), recording location, and their interaction. (D) Accuracy of decoding from cells’ activity shown as the difference from random predictions based on reward zone occupancy probability. Decoders evaluated on the training dataset. ***p < 0.001.

